# Cell-free soluble expression of the membrane protein PsbS

**DOI:** 10.1101/456087

**Authors:** M. Krishnan, T.J.J.F. de Leeuw, A. Pandit

## Abstract

Photosystem II subunit S (PsbS) is a membrane protein that plays an exclusive role in non-photochemical quenching for photoprotection of plants under high-light conditions. The activation mechanism of PsbS and its pH-induced conformational changes are currently unknown. For structural investigation of PsbS, effective synthesis of PsbS with selective isotope or electron-spin labels or non-natural amino acids incorporated would be a great asset. This communication presents cell-free expression as a successful method for *in vitro* production of PsbS that would allow such incorporation. We have optimized the cell-free method to yield soluble PsbS of ~500 ng/µl using a continuous-exchange method at 30°C, along with a successful purification and refolding of PsbS in n-Dodecyl *β*-D-maltoside (*β*-DM) detergent. We expect that the presented protocols are transferrable for *in vitro* expression of other membrane proteins of the Light-Harvesting Complex family.

## Introduction

The discovery of Photosystem II subunit S (PsbS) has revealed that it has a prominent role in sensing changes in thylakoidal pH. PsbS brings about structural rearrangements in the neighbouring photosynthetic proteins, leading to de-excitation of the antenna chlorophylls (Chls), which is known as the fast qE phase of non-photochemical quenching (NPQ) (Li et al. 2000; Li et al. 2004). PsbS is a 22KDa membrane protein with four transmembrane helices and belongs to the light harvesting complex (LHC) protein superfamily (Funk et al. 1995). PsbS can be overexpressed using *E. coli* and refolded into helical structures using several types of detergents (Wilk et al. 2013; Krishnan et al. 2017). Although this overexpression method is well established, there are challenges of inclusion bodies production, low yield while isotope and selective labelling, losses upon insertion into liposomes or nanodiscs and toxic effects to the host cells upon overexpression. Cell-free (CF) protein expression has emerged as an alternative technique for production of a diverse range of membrane proteins for functional studies (Reckel et al. 2008; Schwarz et al. 2008). *In-vitro* refolding of various membrane proteins during the CF reaction can be achieved by adding detergents, liposomes or lipid nanodiscs to the reaction mixture (Ishihara et al. 2005; Schwarz et al. 2008; Katzen et al. 2009). However, the yields for membrane proteins produced using the CF technique is still far below the yields achieved for soluble proteins (Liguori et al. 2007). We show that PsbS from *Physcomitrella patens* can be synthesised and successfully refolded by using a commercial CF system. Various detergents and lipid nanodiscs were added to the reaction mixtures to achieve soluble PsbS using fed-batch system. Purification and refolding of both pellet and soluble PsbS produced using CF reactions was successfully achieved in the detergent n-Dodecyl *β*-D-maltoside (*β*-DM).

## Materials and Methods

### Cell-free (CF) expression protocol

The PsbS gene from *Physcomitrella patens* (Krishnan et al. 2017) was inserted in a pExp5 vector for the CF reactions. Expressway™ Cell-Free Expression System (Invitrogen) was used to carry out the CF reactions We used 25 µl reaction mixture (RM) containing the template DNA plasmid, incubated for 2-4 hours at 30°C or 37°C unless states otherwise, while shaking at 1200 rpm. After 30 minutes, 25 µl of feeding mixture (FM) was added into the reaction volume. To produce soluble PsbS, CF reactions were carried out in the presence of detergents, or in the presence of liposomes or lipid nanodiscs (preparation protocols according to (Crisafi and Pandit 2017)). In the batch feeding method, 12.5 µl of FM was added every 30 minutes for up to 7 hours. In the continuous exchange (CE) method, 50µl of RM and 1 ml FM diluted in HEPES buffer was used for dialysis in 0.1ml 96-well microdialysis plates (Thermo Scientific-Pierce) containing a 10kDa cut-off membrane.

### SDS-page gel analysis

SDS-page gel electrophoresis analysis (12.5% running gel, 4% stacking gel stained with Coomassie brilliant blue R-250 Bio-Rad) was carried out to check the yield of PsbS synthesis along with 2.5 µl of Precision Plus Protein™ Dual Color Standard from Sigma and BSA standard protein.

### Purification of CF-produced PsbS from pellet

CF production of PsbS in the absence of detergents results in pellet containing PsbS. This pellet, P, was subjected to urea wash protocol (Krishnan et al. 2017) to get rid of the impurities.

### Purification of CF-produced soluble PsbS

Soluble PsbS was purified after completion of the CF reaction in the presence of 1%Brij or lipid nanodiscs using Ni-NTA Agarose beads (QIAGEN). The soluble fraction was diluted in equal amounts of binding buffer (50 mM NaH_2_PO_4_, 300mM NaCl, 10mM imidazole and appropriate detergent) and added to the resin, followed by 4-hour incubation at 4°C. The supernatant was removed from the resin by centrifuging for 1 minute at 200xg 4°C after the incubation step. Washing of the resin was carried out three times using 3 volumes of wash buffer (binding buffer + 20mM imidazole). The elution step was carried out 3 times using 1.5 volumes of elution buffer (binding buffer + 250mM imidazole) to achieve purified PsbS.

### Circular Dichroism (CD) spectroscopy

A Jasco J-815 spectropolarimeter (Jasco Labortechnic, Germany) was used to perform CD spectroscopy to check the refolding of the CF-synthesized PsbS. The purified PsbS was buffer exchanged to 50mM mono-sodium phosphate, pH 8.0 with appropriate detergent to remove the high salts and imidazole or urea concentrations. The spectra were recorded between 190nm and 260nm with a scanning rate of 100nm/min, a response time of 8 seconds and an optical path length of 1mm.

## Results and discussion

### CF synthesis of PsbS as pellet

For carrying out the CF reaction, the PsbS plasmid was added to the CF *E*. *coli* reaction mix and incubated at 37°C for 3 hours. Using 1330 ng of plasmid in the CF reaction, a yield of 200 ng/µl PsbS could be produced with this standard reaction protocol (Fig. S1). Since PsbS is a membrane protein, without the presence of detergents, the produced protein formed a pellet, P (Fig. 1a) in the reaction. Fig. 1b shows the purification of PsbS by washing with urea buffer (8M urea,100mM Sodium Phosphate, 100mM Tris) to separate impurities (SS1 and SS2) from the pellet. Upon addition of 0.05% LDS, most of the PsbS from pellet was solubilized (SS3). Further addition of 0.5% LDS to the remaining pellet solubilized the remaining PsbS (SS4). The soluble fraction SS4 was buffer exchanged to remove the urea and stored in the buffer containing 0.1% LDS for further use. Refolding of PsbS from SS4 was carried out in the presence of 0.12% *β*-DM according to a published protocol and analysis of the refolded protein is discussed below (Wilk et al. 2013; Krishnan et al. 2017).

**Fig 1.**
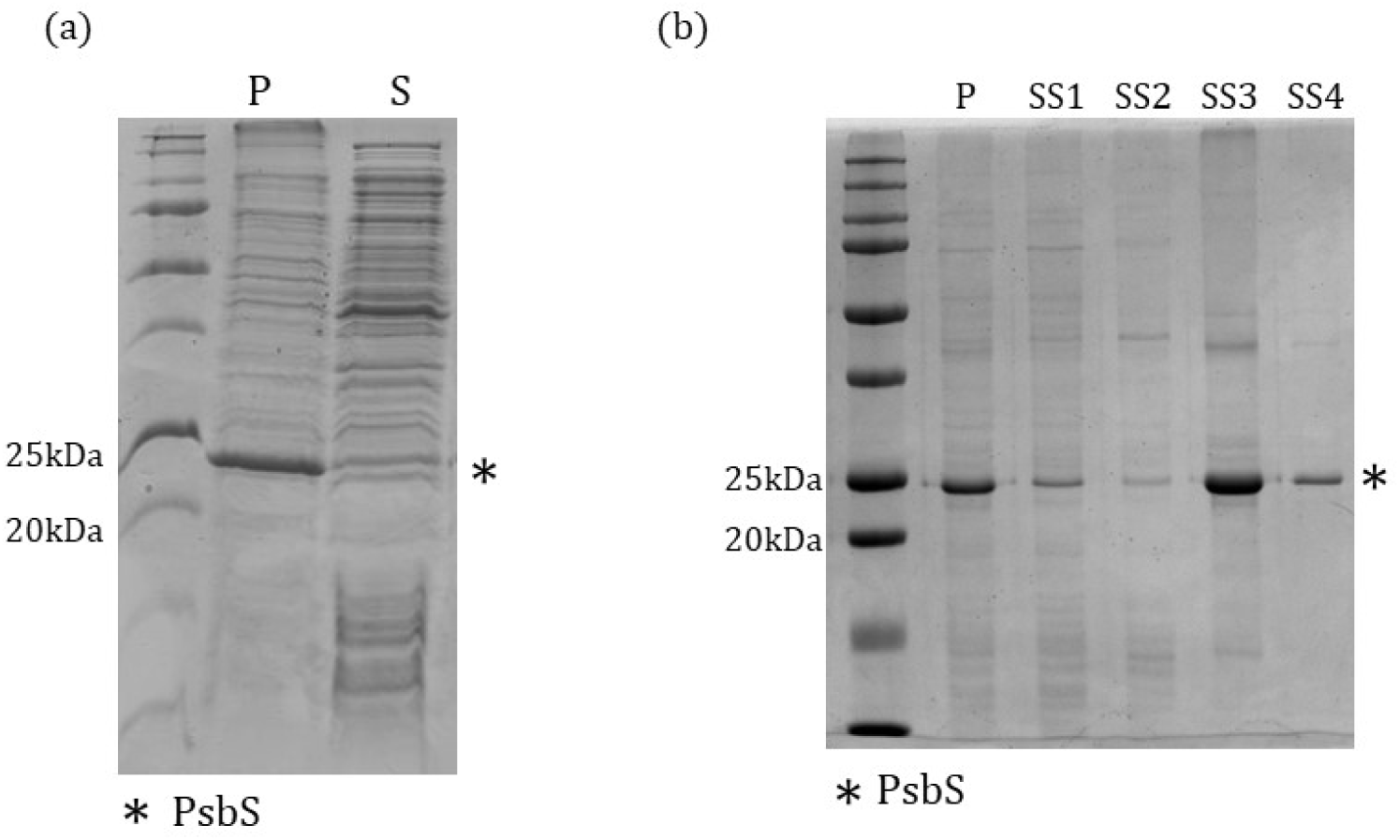
(a) CF production of PsbS (*) in the absence of detergents. CF reactions were separated in pellet (P), and supernatant (S). (b) Purification of CF PsbS from pellet: SS1 and SS2 are soluble fraction after washing the pellet P with 8M urea buffer, SS3 is the soluble fraction after washing with 8M urea buffer and 0.05% LDS and SS4 is the soluble fraction after washing with 8M urea buffer and 0.5% LDS.

### CF synthesis of soluble PsbS

In order to solubilize synthesized PsbS during the CF reaction, the effect of addition of various detergents to the reaction mix was analysed. We tested PsbS solubilization by carrying out the CF reaction in the presence of the detergents *β*-DM, Triton X-100, Brij-35 and Brij-78 (Fig. S2). Furthermore, addition of liposomes or lipid nanodiscs was tested as a method to directly fold synthesized PsbS into lipid membranes (Fig. S3 and Fig. S4). The results are summarized in Table 1 and show that successful production of PsbS was observed in the presence of all the tested detergents and in the presence of liposomes or lipid nanodiscs. In the presence of detergents, the produced protein still precipitated as a pellet, except for reactions carried out in the presence of sufficient amounts of polyoxyethylene-alkyl-ether detergents, Brij-35 and Brij-78. The flexible long chains of polyoxyethylene-alkyl-ethers and their relatively small hydrophilic head might be favouring the solubilisation of membrane proteins (Grip 1982; Garavito and Ferguson-Miller 2001; Külheim et al. 2002). Although some research studies showed that addition of liposomes was beneficial in improving membrane protein yield from CF reactions (Klammt et al. 2005), in our case the presence of asolectin liposomes lowered the production of PsbS considerably.

**Table 1:**
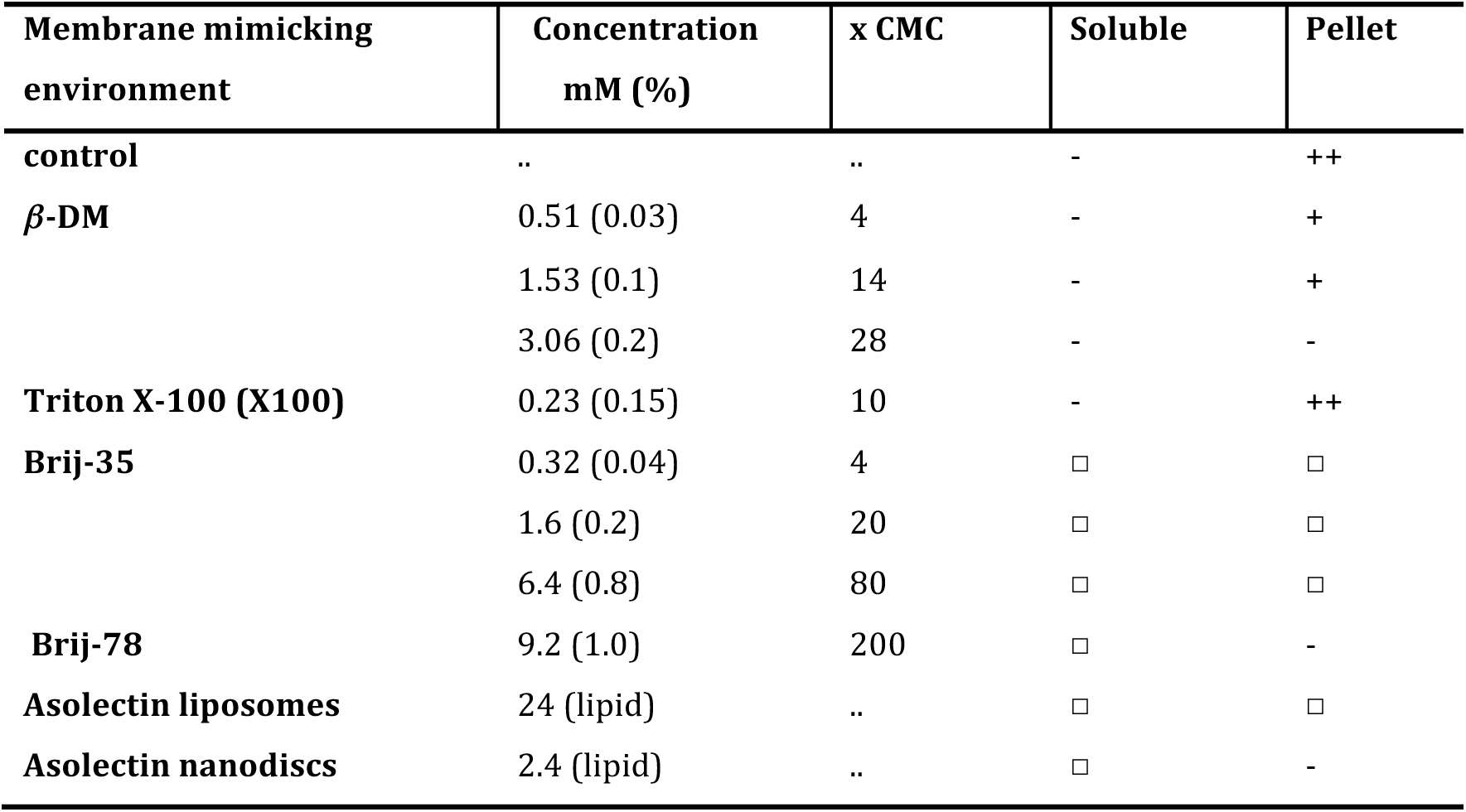
CF expression of PsbS in membrane-mimicking environments. The reaction was carried out at 37°C and PsbS yield as soluble or pellet fraction was classified into four groups: -, no detectable expression; □, spurious expression < 50 ng/µl; +, 51–200 ng/µl; ++, > 200 ng/µl. SDS-page gels of the different reactions are presented in Fig. S1-S4 in the SI section.

### Incubation temperature

The CF reactions were further optimized by varying the incubation temperature, testing reaction temperatures of 37°C, 30°C and 25°C. Since Brij-78 was optimal for expression of soluble PsbS, this detergent was used in the temperature variance experiments. As shown in Fig. S5 and Table 2, reactions carried out at 30°C and 25°C gave a higher yield of PsbS than reactions at 37°C, suggesting that lower temperatures are better suited, which could be due to a slower rate of protein translation. Because part of the produced PsbS was precipitated at 25°C, for further experiments 30°C was selected as the optimal temperature for PsbS production in soluble form.

**Table 2:**
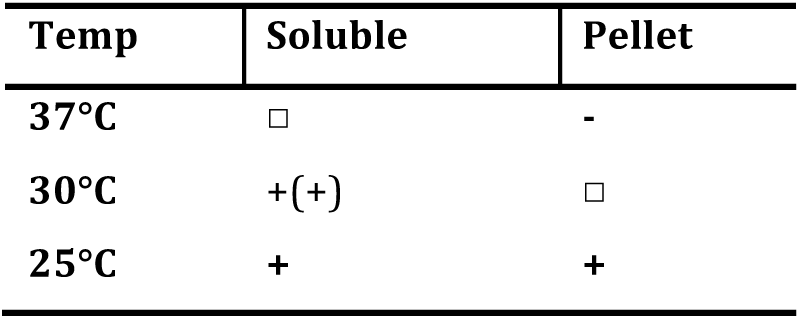
Temperature-dependent CF expression of PsbS. PsbS yield as soluble or pellet fraction was classified into four groups: -, no detectable expression; □, spurious expression < 50 ng/µl; +, 51–200 ng/µl; ++, > 200 ng/µl.

### Batch feeding

During the CF transcription and translation reactions, conditions change because of the consumption of substrates and accumulation of products and by-products, which inhibit the reaction itself. To upscale the PsbS production, batch feeding was tested. In this procedure, instead of adding 25 µl of FM (feeding mix) after 30 minutes, 12.5 µl FM was added up to 10 times every 30 minutes after the reaction was started and the reaction was prolonged up to 8 hours. With this procedure, an amount of 23 µg PsbS could be produced (Fig. 2). The total amount of protein was increased by batch feeding, but also required substantial more FM. Besides that, after 3 hours reaction time, white flakes were formed, suggesting that a part of PsbS was precipitated. The PsbS produced by CF batch feeding reactions in the presence of Brij-78 was purified for analysis (Fig. 2). In addition, we purified PsbS produced in lipid nanodiscs (Fig. S6).

**Fig 2.**
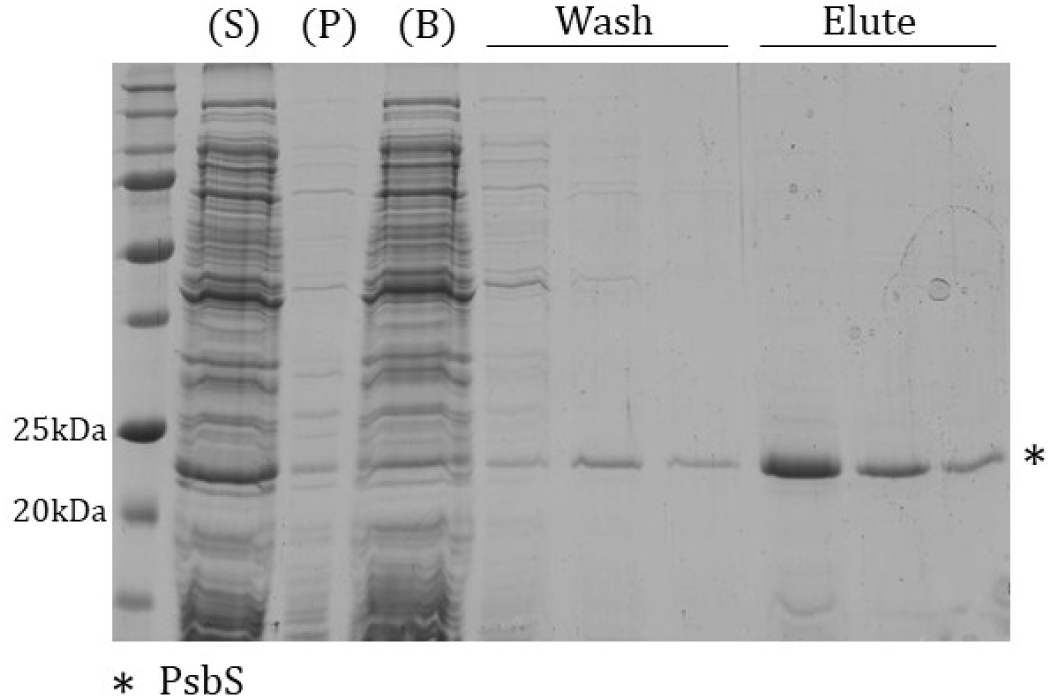
PsbS (*) was expressed at 30°C in the presence of 1% Brij-78 using the batch feeding method; P, pellet and S, supernatant of the reaction mix before loading with Ni-NTA beads. B is the supernatant that was removed after incubation with Ni-NTA beads. Comparison of S and B shows a that most of the soluble PsbS binds to the beads. The beads were washed with buffer containing 20mM imidazole and PsbS was eluted with buffer containing 250 mM of imidazole.

### Continuous exchange

In order to avoiding precipitation during batch feeding and to improve the yield of CF production, we explored the use of continuous-exchange (CE) reactions, where dialysis is used to dilute the inhibitory products that are formed during the reaction in the FM when fresh amino acids and co-factors are added. The CE reaction was carried out using 1% Brij-78 at 30°C for 9 hours with addition of 12.5 µl FM at three time-intervals. Using this set-up, the yield of PsbS was not significantly higher than with the batch-feeding method, but the PsbS did remain in a soluble state. The results are shown in Fig. S7. The yield of PsbS in this reaction was estimated to be ~500 ng/µl and could be further improved by further increasing the reaction time.

### CD spectroscopy of refolded CF-produced PsbS

To check the fold of the CF produced soluble PsbS, UV-CD spectroscopy was carried out by pooling the eluted fractions of PsbS shown in Fig. 2. The initial CD spectrum of the CF-produced PsbS in Brij-78 suggests that the soluble PsbS is in a partly folded molten-globule state and does not show the characteristics of α-helix structure (Fig. S8). To overcome this issue, we first unfolded the PsbS by removal of Brij-78 in the presence of 0.1% LDS followed by refolding in the presence of 0.12% *β*-DM using the standard refolding protocol (Wilk et al. 2013; Krishnan et al. 2017). In addition, PsbS pellets produced from the CF reaction carried out in the absence of detergent (Fig. 1) were refolded using the standard protocol. The CD spectra of CF-produced, refolded PsbS are shown in Fig. 3. According to the CD spectral analysis, the refolded PsbS has ~50% helical structure (Table 3). This number is in agreement with the crystal structure of *spinach* PsbS that shows that PsbS is ~48% α-helical, with the remaining part random coil and a small contribution of *β*-sheet (Fan et al. 2015), and is consistent with CD analyses of detergent-refolded PsbS produced from *E*. *coli* (Krishnan et al. 2017). Comparing the CD spectrum of CF-produced PsbS in *β*-DM to the CD spectrum of *E*. *coli* produced PsbS in *β*-DM, different amplitudes for the characteristic bands at 210 nm and 222 nm are observed. In previous work, we demonstrated that the PsbS CD signature varies for different detergent types, while the estimated helical content is similar. The variations in the CD spectra were attributed to differences in the stromal loop and tail structures of which the folds may depend on the detergent micro-environment. The CD spectra of *E*. coli and CF produced PsbS that are compared in Fig. 3 however are both taken from samples where PsbS is solubilized in *β*-DM. The observed differences could originate from the presence of lipids from the *E. coli* host system in the *E*. *coli* produced PsbS as NMR spectra show the presence of protein-associated lipids that are purified along with the protein (*data not shown*). We suspect that lipids mediate the refolding of PsbS and influence the conformations of the non-helical contents. The influence of lipids on the refolding of membrane proteins is an interesting aspect, which is not easily controlled in recombinant expression systems using host cells, but which could be further explored in CF synthesis, where selective lipids can be added to the synthesis reaction or during the subsequent refolding steps.

**Fig 3.**
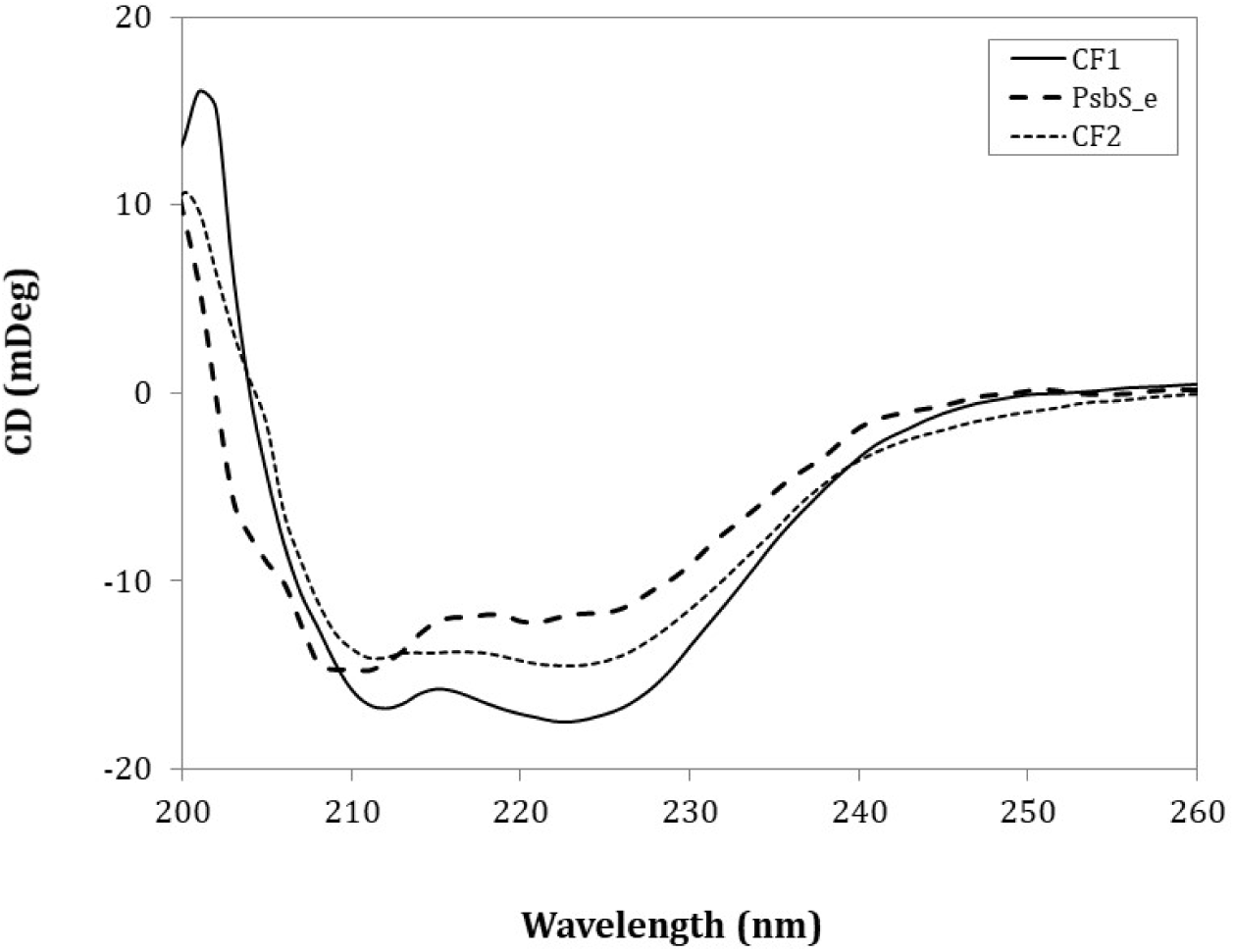
CD spectra of cell free produced PsbS in comparison with *E. coli* produced PsbS refolded in 0.12% *β*-DM. CF1 is PsbS produced with Brij-78 and refolded in *β*-DM. CF2 was produced without a detergent as a pellet, purified and refolded in *β*-DM, PsbS_e was produced from *E. coli*, purified and refolded in *β*-DM.

**Table 3.**
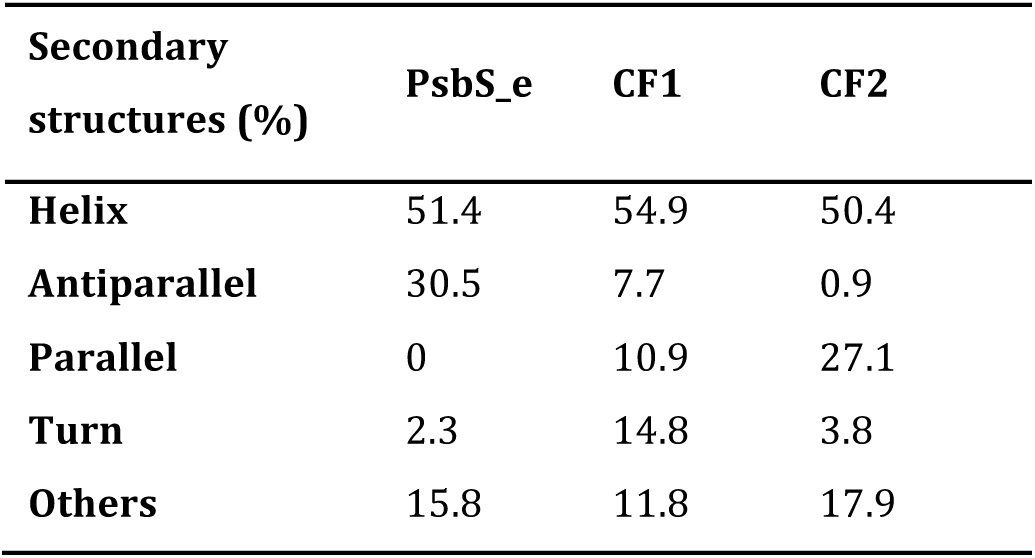
Estimation of secondary structures using BeStSel (Beta Structure Selection (Micsonai et al. 2015)).

| Secondary structures (%) | PsbS_e | CF1 | CF2 |
| --- | --- | --- | --- |
| **Helix** | 51.4 | 54.9 | 50.4 |
| **Antiparallel** | 30.5 | 7.7 | 0.9 |
| **Parallel** | 0 | 10.9 | 27.1 |
| **Turn** | 2.3 | 14.8 | 3.8 |
| **Others** | 15.8 | 11.8 | 17.9 |

## Conclusions

We demonstrate that with CF synthesis, PsbS can be produced either as aggregate pellet or in a soluble state in the presence of membrane-mimicking additives, in contrast to the *E. coli* overexpression, where PsbS is always secreted in inclusion bodies. The CF produced PsbS protein could successfully be refolded into detergent micelles, showing ~50% helical content. The protocols were optimized to yield ~500 ng/µl PsbS production in a single reaction, which could be upscaled for structural studies to produce milligram amounts of protein.

## Acknowledgements

We would like to thank Nora Goosen and Geri Moolenaar for their technical help and advice and Emanuela Crisafi for providing the liposomes. A.P. and M.K. were financially supported by a CW-VIDI grant of the Netherlands Organization of Scientific Research (grant nr. 723.012.103).

